# Gene expression in the hypothalamus of gonad-sex chromosome divergent embryonic chickens

**DOI:** 10.64898/2026.11.15.766747

**Authors:** Vincent Mullis, Jérôme P. Plumier, Zeyu B. Yang, Kiran K. Soma, Nicole M. Baran

## Abstract

Sexual differentiation of the brain of birds is shaped by gonadal steroid hormones and also by cell-autonomous mechanisms resulting from limited sex chromosome dosage compensation. In this study, we compare gene expression in the hypothalamus of gonad-sex chromosome divergent embryonic chickens treated with the aromatase inhibitor fadrozole to their control counterparts. Consistent with a lack of dosage compensation, we find a robust effect of sex chromosomes on hypothalamic gene expression, and a large portion of differentially expressed genes were on the Z and W sex chromosomes. Despite the presence of developing testes and markedly reduced estrogen concentrations in fadrozole-treated chromosomal female (ZW) birds, we find modest effects of gonadal hormones on gene expression in the brain. Weighted gene coexpression network analysis revealed that the most prominent effects were attributable to interactions between chromosomal sex and treatment. Our results highlight that chromosomal sex and steroid hormones together shape the sexual differentiation of the embryonic avian brain, challenging the primacy of the organizational effects of sex steroids on the brain during development.

## Introduction

In all vertebrates, sexual differentiation of the brain and other tissues is shaped by a complex interplay of genetics, hormones, and the environment during development ^1^. In species with sex chromosomes, such as mammals and birds, genes on these chromosomes drive the process of sexual differentiation by initiating heteromorphic gonadal development, which in turn influences the production of steroid hormones that affect gene expression throughout the body ^2^. Note that we intentionally use terminology throughout to describe variation in sex-related phenotypes in a way that is operational, precise, and inclusive ^3,4^.

In mammals, the Sex-Determining Region Y (*SRY*) gene on the Y chromosome initiates the development of the testes from relatively undifferentiated gonads, and testicular cells then produce anti-Müllerian hormone (AMH) and the steroid hormone testosterone ^5,6^. Testosterone (T) and other androgens play a central role in male-typical development of the accessory reproductive structures, the brain, and other tissues. Notably, many of the permanent, organizational effects of T in the brain are mediated by its conversion to 17β-estradiol (E_2_) via the enzyme aromatase and subsequent activation of estrogen receptors ^7,8^.

Sexual differentiation in birds is different from that in mammals in several important ways. First, birds have an independently evolved ZZ/ZW chromosomal system and differentiation of the multipotential gonads into testes is initiated by a double dosage of the Z-linked gene *doublesex and mab-3 related transcription factor 1* (*DMRT1*) gene in ZZ (chromosomal male) individuals ^9–12^. If a single Z-chromosome is present, as it is in ZW (chromosomal female) individuals, a single ovary will typically develop. *DMRT1* inhibits the transcription factor *FOXL2*, which drives expression of *CYP19A1* (cytochrome P450 aromatase) ^13^. E_2_ plays a central role in ovarian differentiation in birds and, unlike in mammals, both P450 aromatase and E_2_ are present in ovaries prior to gonadal differentiation ^14,15^. Thus, a double dose of *DMRT1* inhibits ovarian development and initiates testis development, whereas a single dose of *DMRT1* leads to estrogen synthesis and ovarian development. Gonadal development in birds can thus be altered via manipulations of E_2_. Aromatase inhibitor treatment prior to gonadal differentiation causes ZW embryos to develop testes permanently ^16^ and E_2_ treatment temporarily leads to ovarian-like development in ZZ individuals ^17^.

Second, birds appear to lack a global mechanism for gene dosage compensation. In mammals, XX (chromosomal female) individuals show near complete inactivation of one of their X chromosomes ^18–22^. This is related to a third unique aspect of avian sexual differentiation: the strong role of cell-autonomous sex identity (CASI). CASI is the differentiation of individual cells based on what sex chromosomes are present in their nuclei, independent of gonads and sex hormone concentrations. While CASI occurs in mammals ^23,24^, the lack of gene dosage compensation in birds means that CASI exerts greater influence over sexual differentiation of birds ^25^. Indeed, genes on the Z chromosome have, on average, twice the expression in males as compared to females ^18,19^.

The importance of CASI is especially clear in gonad-sex chromosome divergent birds ^26,27^. In previous studies in which testes were induced during embryonic development in ZW individuals (i.e. ‘sex reversal’), the birds’ appearance and behaviors were not fully male-typical in adulthood ^28^, consistent with the idea that many traits were not shaped by gonads or gonadal hormones, but by the sex chromosomes in specific cells. Similarly, gynandromorph birds, which are bilateral chimeras composed of ZZ and ZW cells, can have either testes or ovaries (or both), depending on the relative sex chromosome composition ^29–31^. Gynandromorphs often exhibit striking phenotypes: male-typical structures and plumage on one side of the body, including sexually heteromorphic structures in the brain, and female features on the other side ^25,29–32^. Additionally, *DMRT1*-knockout ZZ chickens (with ovaries instead of testes) retain many aspects of male-typical appearance ^33^.

Sex steroid hormones produced by the developing gonads play a critical organizational role in the brain during development in birds, like in mammals, but with additional complexity. In several avian species, female-typical mating behavior is induced by exposure to E_2_ *in ovo* in ZZ birds ^34–36^. In Japanese quail, embryonic exposure to E_2_ alters the size of neurons, aromatase activity, vasotocin innervation, and microglia numbers in the adult medial preoptic area of the hypothalamus, an extremely sexually heteromorphic brain region that regulates mating ^37–43^. This suggests that early actions of steroid hormones initiate changes in gene expression in the brain, leading to these long-term molecular, structural, and connectivity differences.

However, the organizational roles of steroid hormones have remained puzzling in the context of another striking sexually heteromorphic behavior in some birds: singing (see ^43^ for review). Although early life E_2_ exposure induces female-typical mating behaviors in ZZ (chromosomal male) zebra finches, it also induces male-typical singing behavior in ZW (chromosomal female) zebra finches ^44–47^. Multiple investigations failed to find evidence that estrogen levels differ during early development between ZZ and ZW birds, and studies about the specific timing of exposure needed to induce these changes produced conflicting results ^48–52^. Recent findings from gynandromorph finches and transcriptomic experiments following treatment during development with an aromatase inhibitor point towards a complex and important role for CASI in combination with both organizational and activational effects of steroid hormones ^29,53,54^. Each brain region is likely differentially impacted by early hormone exposure, suggesting that CASI and the organizational effects of hormones are perhaps more complex in birds than in mammals.

Mating behavior and singing are among the most sexually heteromorphic behaviors regulated by some of the most heteromorphic brain structures identified in the animal kingdom, and nonetheless the mechanisms underlying the differentiation of these structures have remained elusive. Indeed, there is only limited evidence from any avian species about how steroid hormones (particularly estrogens), sex chromosomes, and their interactions influence gene expression during the early sexual differentiation of the avian brain.

Here we aim to disentangle the effects of chromosomal sex and sex steroid hormones on the development of a brain region critical for mating. To do this, we created gonad-sex chromosome divergent embryonic chickens (*Gallus gallus domesticus*) via treatment on day four of incubation with an aromatase inhibitor. This treatment created gonad-sex chromosome divergent ZW (chromosomal female) embryos that developed testes ^55^. We dissected treated and control ZZ and ZW embryos after seven additional days of incubation and collected plasma samples to quantify steroid hormones using liquid chromatography-tandem mass spectrometry (LC-MS/MS), blood samples for chromosomal sexing, whole frozen heads for brain tissue collection, and visually phenotyped the embryonic gonads. We then performed RNA-sequencing (RNA-seq) on tissue punches from the developing hypothalamus and carried out differential expression analysis.

Our results reveal an outsized role for CASI in the development of the avian brain, alongside complex interactions between sex chromosomes and gonadal hormones. We observed a more moderate effect of hormonal sex at this early stage of neurodevelopment and we found evidence that steroid hormones impact multiple cell types and functional processes in the developing brain, including astrocyte function, mitochondrial metabolism, and neuronal development. Our findings suggest that organizational effects of hormones are less widespread in birds than in mammals and demonstrates a tractable approach to understand how steroid hormones and sex chromosomes interact.

## Results

### 2.1 Gonad/chromosome comparison between sample groups

To measure the differing impacts of sex chromosomes and sex hormones, we injected 63 four-day-old chicken eggs with either 0.1 mL of sterile physiological saline (control) or 1mg of fadrozole (F3806 from Sigma-Aldrich, Saint-Louis, MO) dissolved in 0.1 mL of saline (Figure 1A & B). Of these eggs, 52 (n = 26 saline, n = 26 fadrozole) were able to be phenotyped at E11. Of the saline-treated eggs, 18 developed ovaries and 8 developed testes, while of the fadrozole-treated eggs, 6 developed ovotestes (a gonad that was ambiguous or in between typical ovaries and testes in the degree of asymmetry) and 20 developed testes. Genotyping revealed that all embryos treated with saline had gonads aligning with chromosomal sex, while in those treated with fadrozole, all ZZ embryos developed testes and all ZW embryos developed testes or ovotestes (Figure 1D).

**Figure 1.**
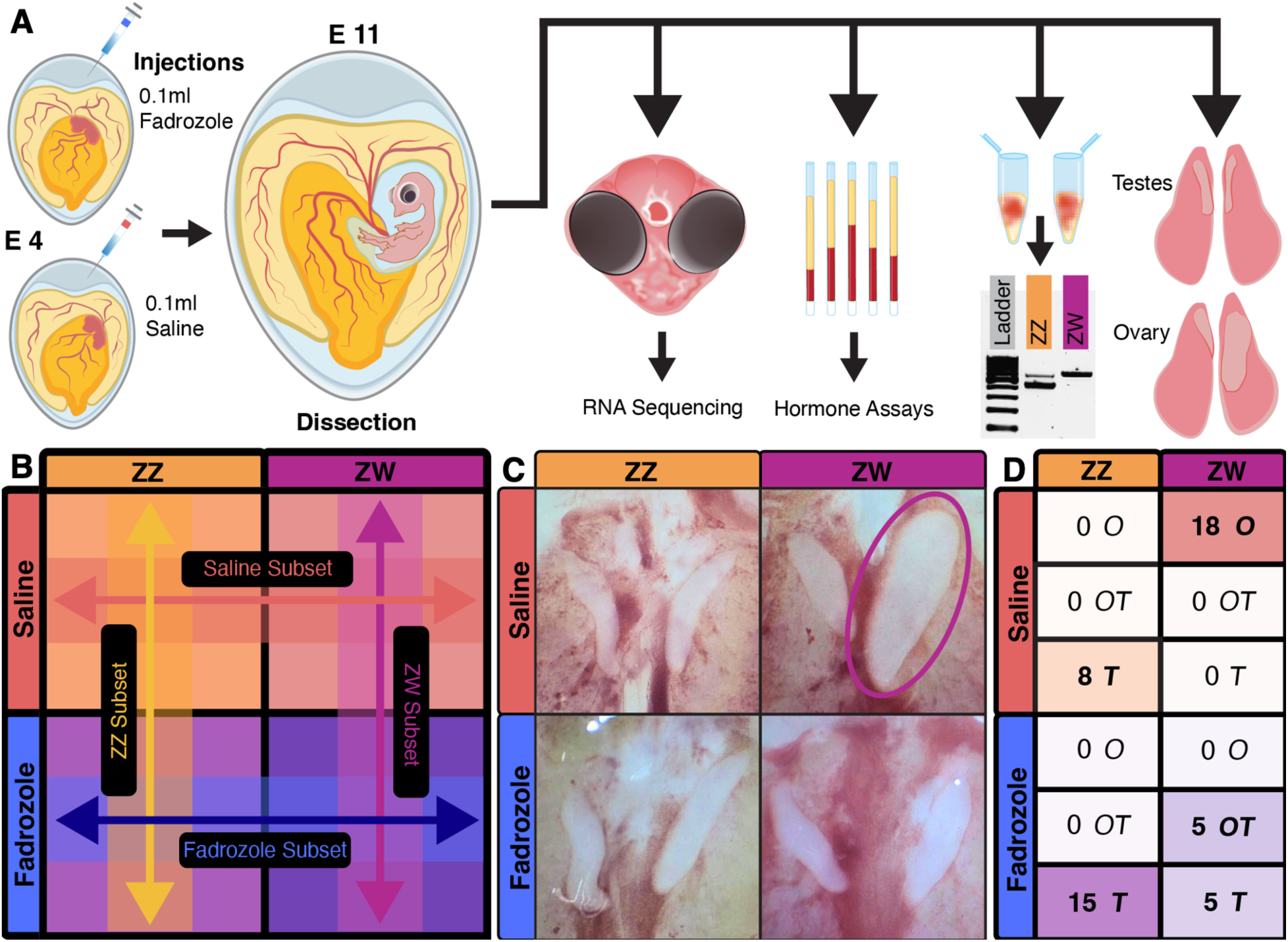
Experimental design and results. **A)** Schematic of experimental procedures. Fertilized eggs were injected with saline (control) or fadrozole on day 4 of incubation (E 4). During dissection on day 11 (E 11), gonads were phenotyped, whole heads were collected for RNA-seq, plasma samples were collected for hormone concentrations, and blood/tissue samples were collected for chromosomal sexing. **B)** The 2×2 experimental design showing contrasts between the four sample groups. **C)** Gonad phenotype examples from each of four sample groups, the single ovary in the control ZW group is circled. Images shown as pictured through the microscope, so the anatomical left is on the right side of the image. **D)** Table summarizing the resulting sample gonadal phenotype counts: ovary (O), ovotestes (OT), and testes (T).

We chose 20 samples (5 per treatment and sex) for RNA-seq and hormone measurement based on the quality of gonadotyping, genotyping, tissue punch, and plasma sample. For one of these samples, the PCR result was incorrect and it was later discovered to have ZW instead of ZZ chromosomes upon examining the RNA-seq data. This meant the sample distribution across the four sample groups included: *n* = 4 ZZ control, *n* = 6 ZW control, *n* = 5 ZZ fadrozole-treated, and *n* = 5 ZW fadrozole-treated samples.

### 2.2 Plasma hormone analyses

For the 20 plasma samples, we quantified a panel of steroid hormones using LC-MS/MS. Our results confirmed that the fadrozole treatment worked as expected, as both E_2_ (Fig 2A) and estrone (Fig 2B) concentrations were decreased to nearly non-detectable levels in fadrozole-treated ZW (chromosomal female) birds. Indeed, using aligned rank transformed ANOVA followed by aligned rank transform contrasts ^56^, we found that there were significant main effects of treatment (*F*(1,16) = 42.35, *p* < 0.001) and genotype (*F*(1,16) = 50.29, *p* < 0.001), and a significant treatment:genotype interaction (*F*(1,16) = 24.80, *p* < 0.001) for E_2_. The saline-treated ZW birds showed higher E_2_ than the other groups, with all three other groups being statistically indistinguishable from each other. A similar pattern was observed for estrone: we found significant main effects of treatment (*F*(1,16) = 42.20, *p* < 0.001) and genotype (*F*(1,16) = 79.41, *p* < 0.001), and a significant treatment:genotype interaction (*F*(1,16) = 7.20, *p* = 0.016).

**Figure 2.**
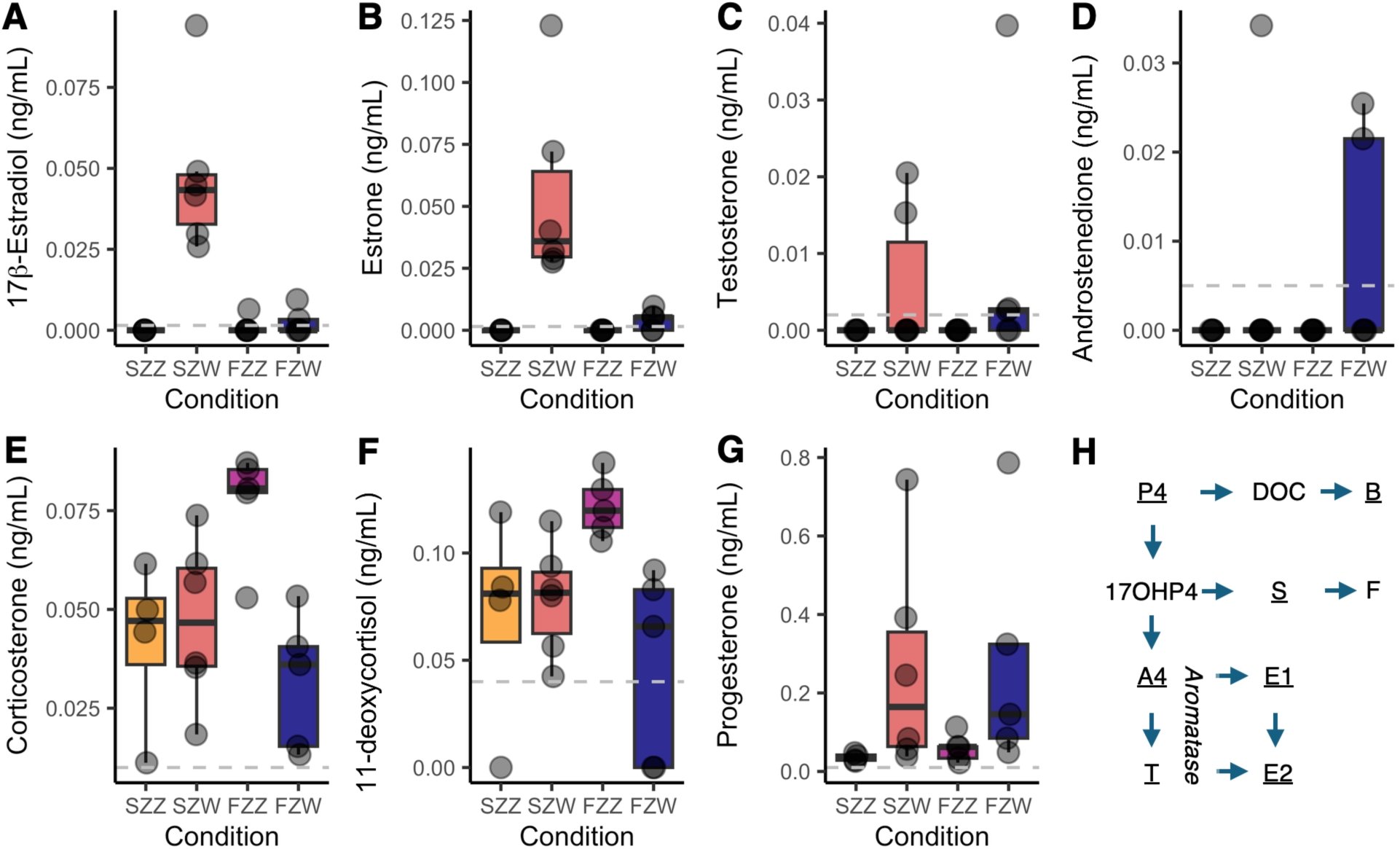
Boxplots of steroid hormone measurements obtained from liquid chromatography tandem mass spectrometry (LC-MS/MS) in plasma. Individual data points are plotted in transparent light gray. Plasma was collected from embryonic chicken trunk blood at day 11 of incubation for **A)** 17β-estradiol, **B)** estrone, **C)** testosterone, **D)** androstenedione, **E)** corticosterone, **F)** 11-deoxycortisol, and **G)** progesterone. The gray dashed lines show the lower limits of quantification (LLOQ). All samples with measurements below the LLOQ are plotted as zero. In addition, we also examined the following steroids using LC-MS/MS, but the concentrations were below the LLOQ for all samples: androstenediol, cortisol, dehydroepiandrosterone, 5α-dihydrotestosterone, estriol, and pregnenolone. **H)** Illustration of the steroid synthesis pathway using abbreviations. Progesterone, P4; 17*a*-hydroxy progesterone, 17OHP4; androstenedione, A4; testosterone, T; 11-deoxycorticosterone, DOC; 11-deoxycortisol, S; estrone, E1; 17β-estradiol, E_2_; corticosterone, B; cortisol, F. All steroids for which data are shown are underlined. The treatment conditions are: SZZ, saline-treated ZZ; SZW, saline-treated ZW; FZZ, fadrozole-treated ZZ; FZW, fadrozole-treated ZW.

We hypothesized that fadrozole treatment might increase androgens, but we found no evidence of this. Instead, we were surprised to find a significant main effect of genotype only (*F*(1,16) = 10.66, *p* = 0.005), with ZW (chromosomal female) birds having higher T (Fig 2C). Notably, however, only 5/11 ZW birds had detectable T at all. There was also a non-significant trend for androstenedione to be higher in ZW (chromosomal female) birds (*F*(1,16) = 4.00, *p* = 0.062, Fig 2D). Unexpectedly, we found that fadrozole-treated ZZ (chromosomal male) birds had increased glucocorticoid concentrations. For corticosterone, the primary glucocorticoid in birds, there was a significant main effect of genotype (*F*(1,16) = 9.13, *p* = 0.008) and a significant treatment:genotype interaction (*F*(1,16) = 8.17, *p* = 0.011), with fadrozole-treated ZZ birds showing a higher concentration of corticosterone than fadrozole-treated ZW birds (Fig 2E). A similar pattern was found for 11-deoxycortisol, where we found a significant main effect of genotype (*F*(1,16) = 5.69, *p* = 0.030) and a significant treatment:genotype interaction (*F*(1,16) = 5.78, *p* = 0.029, Fig 2F). Interestingly, progesterone was higher in ZW (chromosomal female) birds, where we found a significant main effect of genotype (*F*(1,16) = 8.86, *p* = 0.009, Fig 2G).

Taken together, these results indicate that the aromatase inhibitor fadrozole significantly decreases estrogen synthesis (Fig 2H), without an impact on androgens. Androgens and progesterone are higher in ZW (chromosomal female) birds, regardless of treatment. One unexpected finding emerged suggesting that fadrozole-treated ZZ (chromosomal male) birds exhibited higher glucocorticoid synthesis.

### 2.3 RNA quality control & analysis

To determine differential expression between sample groups, we extracted RNA from tissue punches of the hypothalamus for our 20 samples. RNA-sequencing yielded on average 30 million reads per sample. The average GC content per sample was 46.38%. No sequences across all samples were flagged for poor quality by FastQC and HISAT2 had an overall alignment rate of 93.7% (Supplementary Table 1).

### 2.4 Differential gene expression analysis

Using principal components analysis (PCA), we show that chromosomal sex has a large effect on gene expression, and treatment has a smaller, but prevalent, effect as well (Fig 3A). Principal component 1 (Feature 1, accounting for 58% of the variance between samples) shows clear separation of samples by sex chromosome. Principal component 2 (Feature 2, 16% variance) shows near complete separation by treatment group.

**Figure 3.**
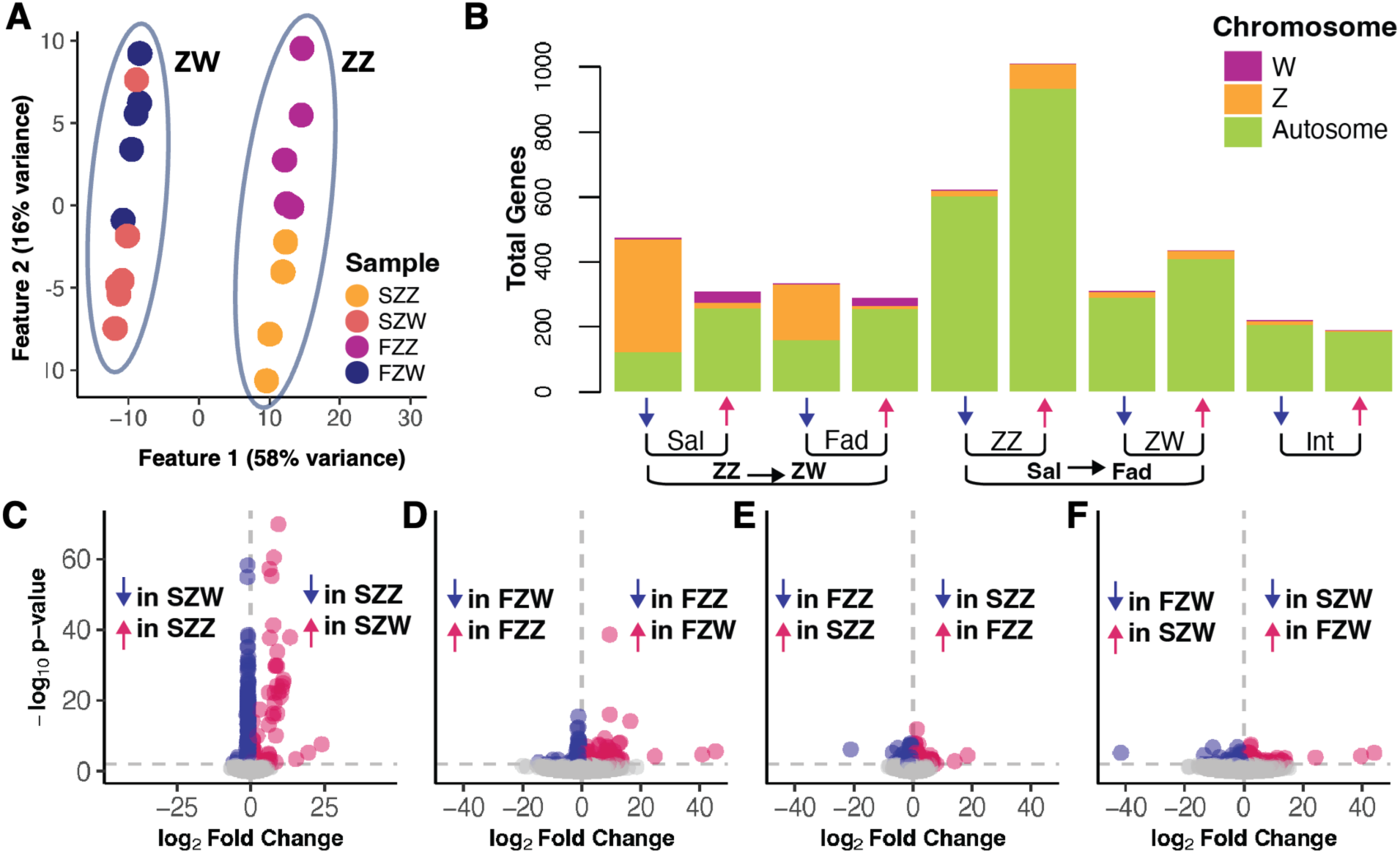
Differential expression analysis results. **A)** A principal component analysis (PCA) plot showing Feature 1 which separates samples by chromosomal sex and accounts for 58% of variance, and Feature 2 which separates largely by treatment group and accounts for 16% of variance between samples. **B)** A barplot showing numbers of differentially expressed genes by chromosome (autosome, Z, or W chromosome) for each DESeq2 contrast when using the full model with interaction term (see STAR Methods). DESeq2 contrasts are linear combinations of estimated log_2_fold changes which test if differences between groups are equal to zero. ZZ→ZW are genes that are either up- or down-regulated in ZW compared to ZZ. Sal→Fad are genes that are either up- or down-regulated in fadrozole treatment compared to saline control. Within these groups, we then consider the effect of treatment or genotype, respectively. Int are genes that show differences in the effect of treatment depending on genotype, i.e. the interaction. **C-F)** Volcano plots showing significant genes (up-regulated in red and down-regulated in blue) plotted according to log_2_ Fold Change (x axis) by -log_10_ p-value (y axis) by **C)** Expression of genes in SZW samples in relation to SZZ sample level expression, **D)** FZW genes expression compared to FZZ, **E)** SZZ genes expression compared to FZZ, **F)** SZW genes expression compared to FZW.

We then used *DESeq2* to identify genes that were differentially expressed between chromosomal sex and treatment groups (see STAR methods for details). When comparing saline-versus fadrozole-treatment ZZ (chromosomal male) samples, we unexpectedly found considerably more differentially expressed (DE, *p_adj_* < 0.01) genes (1635 DE genes) than other contrasts. It was more than twice the number of genes differentially expressed in fadrozole-treated compared to control-treated ZW (chromosomal females) samples (749 DE genes), where fadrozole treatment induced gonad-sex chromosome divergence. There was also a comparatively smaller number of differentially expressed within each treatment when comparing the two chromosomal sexes (SZZ vs SZW, 790 DE genes; FZZ vs FZW, 627 DE genes), and only 416 genes showed differential expression between treatment groups depending on the chromosomal sex (i.e. a chromosome x treatment interaction) (Supplementary Table 2).

When these gene subsets are further divided into up- and down-regulated genes located on sex chromosomes versus autosomes, distinct patterns emerge (Fig 3B). In both contrasts between chromosomal sex groups, we see in ZW individuals a consistent and expected pattern of decreased expression of genes on the Z chromosome and greater expression of genes on the W chromosome compared to ZZ individuals (Fig 3B). In comparison, the majority of genes differentially expressed when comparing treatment groups are autosomal. The SZZ vs FZZ contrast identified slightly more differentially expressed genes on the Z chromosome that were down-regulated compared to other treatment comparisons.

We also observe very low p-values in genes of SZZ vs SZW contrast, reflecting the robust effect of chromosomal sex and gonad-sex chromosome concordance on expression (Fig 3C). In comparison, genes in the FZZ vs FZW contrast exhibit higher p-values, indicating that fadrozole treatment may produce more variability in gene expression or reduced differentiation in gene expression between the chromosomal sexes in the context of fadrozole treatment (Fig 3D). There is a large fold-difference in up-regulated genes when comparing chromosomal sexes, likely reflecting the presence of genes on the W chromosome in ZW individuals and double dosage of many genes in ZZ individuals. While the SZZ vs FZZ and SZW vs FZW contrasts do not have as low of p-values as the chromosomal contrasts, there are still many genes differentially expressed, with some having very high fold differences (Figures 2E-F).

To determine the potential biological functions underlying gene expression differences, we conducted gene ontology (GO) term analysis using the lists of significantly differentially expressed genes for each contrast subdivided by up- and down-regulation. This analysis revealed profiles of expression associated with both chromosomal sex and treatment (Supplementary Table 3 & 4). In both comparisons between chromosomal sex groups (SZZ vs SZW and FZZ vs FZW), there was consistent enrichment of terms relating to gene expression (e.g. transcription and epigenetic regulation) in up-regulated genes and of terms relating to translation machinery in down-regulated genes. In the SZZ vs SZW contrast, up-regulated genes also had enrichment of bioenergetics-related terms, and the down-regulated genes had enrichment in terms relating to cellular components. In the treatment contrasts (SZZ vs FZZ and SZW vs FZW), genes down-regulated with fadrozole treatment were associated with synaptic signaling and neurogenesis. In the genes up-regulated with fadrozole treatment, there was enrichment for terms relating to developmental processes and gene expression.

### 2.5 Weighted gene co-expression network analysis (WGCNA)

To better understand the interactions between chromosomal sex and sex hormones, we performed weighted gene co-expression network analysis (WGCNA). Of the 24 modules identified via WGCNA (Supplementary Table 5), 17 had eigengenes with expression patterns significantly correlated (*p* < 0.01) with at least one of the four main pairwise comparisons between groups (SZZ vs SZW, FZZ vs FZW, SZZ vs FZZ, SZW vs FZW, Supplementary Fig 1 for p-values).

Each module has a module eigengene (ME) representing the overall gene expression patterns of genes within the module. To determine which modules were related to chromosome and treatment conditions, we examined each module’s module-trait associations (MTA) (Fig 4A, Supplementary Table 5).

**Figure 4.**
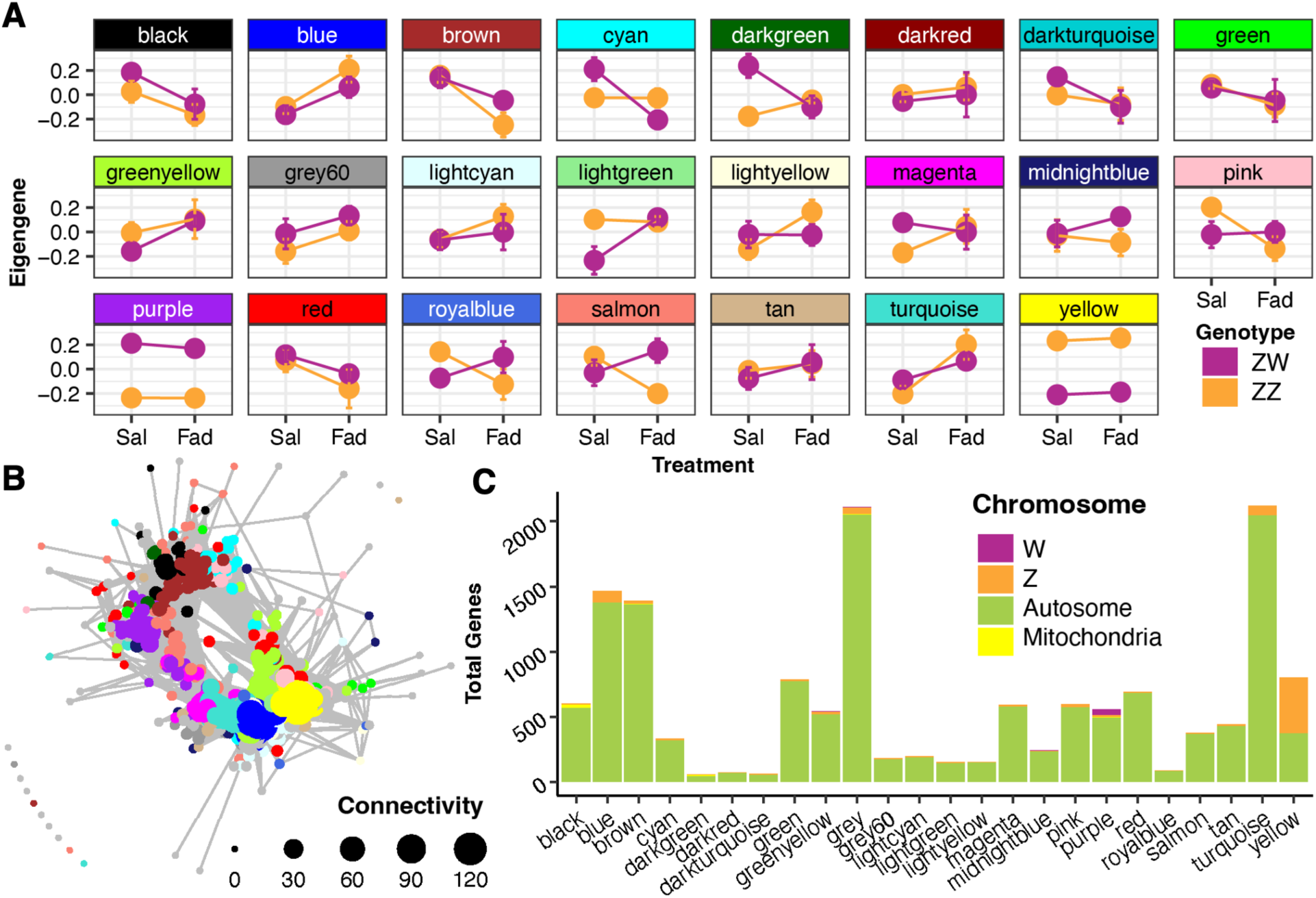
Weighted gene co-expression network analysis results. **A)** Distribution and mean ± SE of module eigengene values across chromosomal sex and treatment groups for each module. Error bars represent standard error of individual samples of the mean eigengene value for each sample group by module. **B)** Co-expression network showing relative relationships between hub genes within modules. **C)** A barplot showing the distribution of each module’s genes across the Z, W, and autosomal chromosomes.

The purple and yellow modules were clearly associated with sex chromosome, with purple having higher expression in the ZW samples (MTA, p < 0.001) and yellow in ZZ samples in both treatment groups (MTA, p < 0.001 for all chromosome contrasts). In addition to this effect of chromosome, there also appears to be a significant effect of treatment in the yellow module, where genes are up-regulated between the control and fadrozole treated ZZ samples (MTA, *p* = 0.01). The blue module shows consistent up-regulation with fadrozole treatment for both chromosomal sexes (Fig 4A, Supplementary Table 5).

Several modules show more complex interactions, where significant effects of treatment differ between chromosomal sex groups. The black, brown, cyan, greenyellow, lightgreen, lightyellow, pink, royalblue, salmon, and turquoise modules, for example, all have one chromosomal sex mostly unaffected by treatment while the other experiences significant upwards or downwards expression with treatment. The darkgreen module has an even more complex expression pattern where chromosomal sex groups respond significantly in opposite manners to treatment. In both of these patterns, the chromosomal sex groups often start at the same level of expression and diverge significantly with treatment, or vice versa.

To understand the relationships between these modules we plotted the top most significant hub genes (adjacency threshold = 0.05) for each module in a co-expression network based on the similarity of their expression patterns (Fig 4B). In this plot, we can see that the purple module (genes higher in ZW samples) clusters with several modules in which expression in ZW samples is down-regulated with fadrozole treatment (black, brown, cyan, darkgreen). The yellow module (genes higher in ZZ samples) forms a second cluster with the blue, greenyellow, grey60, lightcyan, magenta and tan modules, which all see increased expression in ZZ samples with fadrozole treatment. A few modules (green, pink, red and salmon) show hubs spread throughout the plot and all show decreased expression in ZZ samples post fadrozole treatment.

When the genes of each module were mapped to chromosomes, we observed that most modules have low numbers of genes on the sex chromosomes and an overrepresentation of genes on autosomes (Fig 4D, Supplementary Table 6). Exceptions to this are the yellow module, with a drastic overrepresentation of genes on the Z chromosome (*n* = 421*, ꭓ*^2^(1) = 2179.2, *p* < 0.0001), and the purple module, with 44 genes on the W chromosome, representing an overrepresentation from typical gene distribution (*n* = 44*, ꭓ*^2^(1) = 153.3, *p* < 0.0001). The overrepresentation of sex chromosome genes in these modules makes sense, given that they are largely associated with chromosomal sex. There was also a large overrepresentation of mitochondrial genes in the black (*n* = 24, *ꭓ*^2^(1) = 68401.3, *p* < 0.0001) and darkgreen (n = 24, *ꭓ*^2^(1) = 102,069, *p* < 0.0001) modules.

GO term analysis of the modules revealed several strong associations with biological functions, with multiple modules having highly enriched, statistically significant, and thematically consistent GO terms (Supplementary Table 7). These biological themes included synaptic/neuronal signaling and development (black, brown, cyan, and pink), translation and translation machinery (green and royalblue), cell cycle/mitosis (lightyellow, turquoise), mitochondrial activity (red), epigenetic regulation (tan and turquoise), morphogenesis (greenyellow), immunity (blue and greenyellow) and astrocyte function and vasculature (blue). Other modules yielded fewer GO terms but included biological profiles related to intracellular structure (darkgreen), proteostasis (darkturquoise), hormone signaling (grey), cellular adherence (lightgreen), synapses (magenta, salmon), and DNA transcription (salmon). Finally, GO term analysis revealed that many modules did not have any significant GO terms associated with them, including the dark red, grey60, lightcyan, midnight blue, purple, and yellow modules. Overall, however, we see that many modules have gene expression patterns representing functional differences.

### 2.6 Comparative ChIP-Seq analysis

To test the hypothesis that genes identified in our analyses are regulated by androgen receptor (AR) or estrogen receptors (ERα or ERβ), we compared our DE and WGCNA gene lists to genes exhibiting differential AR ^57^, ERα ^58^, ERβ ^59^ binding in chromatin-immunoprecipitation sequencing (ChIP-Seq) studies. We also compared our lists with a mineralocorticoid receptor (MR) and glucocorticoid (GR) receptor ChIP-Seq study ^60^. We used a *χ*^2^ test of independence to compare the expected overlap by chance in the gene lists to the observed number of genes found in both data sets. Examining our DE gene lists, we found an underrepresentation of genes found in the AR ChIP-seq dataset for all contrasts (*χ*^2^(4) = 30.252, *p* < 0.001) (all other ChIP-seq datasets were not significant). This is consistent with low androgen concentrations across all groups, but especially in ZZ (chromosomal male) birds. We also found a statistically significant association with all tested steroid receptor gene lists and all the WGCNA modules: AR (*χ*^2^(23) = 210.7, *p* < 0.001), ERα (*χ*^2^(23) = 120.5, *p* < 0.001), ERβ (*χ*^2^(23) = 22.7, *p* < 0.001), MR (*χ*^2^(23) = 59.4, *p* < 0.001), GR (*χ*^2^(23) = 47.5, *p* < 0.001) lists. For each steroid receptor, we then asked which of the 24 modules contributed 10% or more to the *χ*^2^ value to determine which modules showed the greatest over-or underrepresentation in the steroid receptor ChIP-seq datasets.

In the AR gene list, the brown, green, grey, purple, and red module genes were all underrepresented. The green, grey, red, and tan modules all contributed 10% or more to the *χ*^2^ value for ERα gene lists, with green, grey, and red modules having an underrepresentation and purple overrepresentation of ERα regulated genes. Interestingly, the green and red modules, and the grey module, which is one filled with unassigned genes, had underrepresentation in both AR and ERα datasets, meaning these modules contain fewer genes than expected by chance that appear to be regulated by these receptors. For the ERβ gene list, only the blue module showed evidence of overrepresentation. For MR receptor genes, the blue, lightgreen, magenta, and tan modules all were overrepresented. The lightgreen, pink, and tan modules also had overrepresentation of GR receptor genes.

## Discussion

Previous studies have shown that treatment with the aromatase inhibitor fadrozole produces gonad-sex chromosome divergence in embryonic chickens ^26^. We find this same effect of fadrozole on gonadal development: all chromosomally female (ZW) chickens treated with fadrozole developed testes or ovotestes.

Our treatment effect was confirmed by hormone measurements, where estrogens (E_2_ and estrone) were drastically lower in fadrozole-treated ZW chickens, as expected. Notably, androgen concentrations did not increase with fadrozole treatment in ZZ or ZW birds. We also uncovered that only ZW (chromosomal female) chickens had measurable concentrations of estrogens and androgens. Previous work has shown that plasma steroids, especially E_2_, are higher in ZW (chromosomal female) embryonic chickens, but rarely have ZZ (chromosomal male) steroid concentrations been close to zero ^61–63^. Given the high sensitivity and specificity of the LC-MS/MS assay, this discrepancy may relate to lower sensitivity and specificity of immunoassays used previously, combined with issues related to high lipid content of embryonic plasma ^61^.

Surprisingly, ZZ (chromosomal male) birds treated with fadrozole had higher concentrations of glucocorticoids than control ZZ birds. Fadrozole may have off-target effects on the 11β-hydroxylase and aldosterone synthase enzymes ^64,65^. In ZZ birds, which have almost no androgens to interact with aromatase, fadrozole may have impacted other steroid synthesis enzymes to influence glucocorticoid concentrations. This also may account for our observation of an unexpectedly large number of differentially expressed genes between control- and fadrozole-treated ZZ birds, as increased glucocorticoids may have influenced gene expression. Several WGCNA modules (lightgreen, tan, blue, magenta, and pink modules) had an overrepresentation of genes regulated by glucocorticoid receptors, though this observation requires further exploration.

Taken together, these hormone results suggest that, at least early in development, male-typical development in embryonic chickens is characterized by near total absence of sex steroid hormones. At first glance, this might provide some support for the idea of male-typical development as the ‘default’ pathway and that the presence of estrogens may serve to ‘demasculinize’ the gonads and nervous system (see ^35,43^ for excellent reviews of the history of this hypothesis). However, our gene expression data point to extremely important roles for CASI and interactions between hormones and sex chromosomes. Many genes were found to have low p-values in their differential expression between chromosomal sexes, especially within control-treated groups (see Figure 3C). We also do not observe fully ‘male-typical’ gene expression in fadrozole-treated ZW (chromosomal female) birds, although fadrozole treatment does decrease the number of differentially expressed genes between ZZ and ZW birds. This perhaps reflects an equalizing effect of converging low hormone levels and the presence of developing testes, but gonadal hormones alone clearly cannot account for the vast majority of gene expression differences in the developing hypothalamus.

Despite the greater variability and fewer differentially expressed genes when comparing control-to fadrozole-treated embryos, genes that were found to be differentially expressed between treatments have a very high log-fold change (Figure 3E & F), suggesting that there is a large impact of sex steroids on at least some genes in the brain. In chicken and quail, exposure to E_2_ *in ovo* leads to female-typical mating behavior in ZZ birds ^34,36^. E_2_ binding to ERɑ in the mouse brain coordinates a sexually-biased gene expression program in the brain ^58^, highlighting the widespread impact of estrogens in brain organization. These results, thus, contribute to our understanding of how estrogens directly influence gene expression programs and sexual differentiation in birds.

Several modules emerged as particularly interesting from the WGCNA analysis. The yellow and purple modules were highly correlated with chromosomal sex, and accordingly had drastic overrepresentation of genes on the Z and W chromosomes, respectively. However, GO term analysis yielded no significant enrichment terms for these modules. This suggests that, despite consistent expression patterns within their groups of genes, they do not have an identifiable signature associated with known functions or cellular components. This lack of identification may be affected by the lack of GO terms assigned to the chicken genome in comparison with mammalian species or potential limitations of WGCNA design and module counts, but it also most likely indicates these effects are purely gene dosage related, rather than a functional association between genes that happen to be on the sex chromosomes.

While many other modules show trends of gene expression between sample groups, few of these trends cross the stringent threshold of statistical significance (*p* = 0.01). The blue module is the only module with statistically significant patterns of expression representative of a primary effect of treatment, where both chromosomal sexes show increased expression of genes in this module when treated with fadrozole. Genes in the blue module also showed enrichment in GO terms relating to astrocytes, blood/blood vessels, and immune cells, which is consistent with a rich literature demonstrating an important role for aromatase expression in astrocytes (and other glial cells) as an important part of sexual differentiation of the brain ^66,67^ and in response to neural injury and inflammation ^68,69^. Genes in the blue module show possible associations with regulation by ERβ in our dataset. In mice, astrocytes highly express ERβ and activation of astrocytes via ERβ has an important neuroprotective role in models of aging and stroke ^70,71^. Together, this suggests that genes in the blue module may be associated with altered astrocyte or microglial function via ERβ in response to fadrozole treatment.

Another interesting module that emerged from our analysis was the black module, which showed a decrease in expression in ZW (chromosomal female) embryos with fadrozole treatment, i.e. the group that showed gonad-sex chromosome divergence. This module showed enrichment for GO terms associated with synaptic communication and an extreme overrepresentation of mitochondrial genes. In mammals, estrogens regulate mitochondrial metabolism, which might underlie sex differences in many diseases ^72^. Our results suggest that estrogens regulate mitochondria in birds, as well. Indeed, in chickens and Japanese quail, steroid receptors are found in the muscle mitochondria, which are thought to be important in physiology and thermoregulation ^73^. However, there is much more to learn about what role steroid-sensitive mitochondria play in the brain early in development in any species.

Unsurprisingly, several modules seem to be strongly associated with neuron function, including the black, brown, cyan, and pink modules. All of these modules show a general trend of decreased expression of the module eigengene with fadrozole, suggesting that the growth and function of neurons was consistently impacted by reduced estrogens. In addition, the royalblue, tan, and turquoise modules all show enrichment of GO terms associated with regulation of gene expression, such as RNA processing and epigenetic regulation. The lightyellow module showed enrichment of GO terms associated with mitosis and cell division, an important process in the developing brain as rapid neurogenesis and cell proliferation occurs. Nearly all of these modules show complex interactions between chromosomal sex and fadrozole treatment, suggesting that although the effects of treatment are widespread, chromosomal sex and interactions between chromosomal sex and steroid hormones are extremely important.

Altogether, these results suggest that effects of both chromosomes and steroid hormones are widespread and complex in the sexual differentiation of the brains of chickens. Multiple cell types appear to be impacted by fadrozole treatment, particularly neurons and astrocytes. The foundational developmental processes of gene expression, cell division, and mitochondrial metabolism are also altered by treatment, depending on the chromosomal sex of the embryo. This is consistent with the idea that steroid hormones play an important role in organizing the developing brain in birds. However, the impact of these hormones is likely cell-type specific and dependent on chromosomal sex. Indeed, our results confirm that CASI plays an especially important role in avian sexual differentiation. Future studies would benefit from cellular resolution via single cell transcriptomics to better understand the nature of these complex interactions.

## Conclusions

Our findings support the idea that the organizational effects of steroid hormones in birds, particularly estrogens, are dependent on sex chromosomes and likely cell-type specific. This helps account for the discrepancies in previous studies investigating the organizational effects of steroid hormones in birds, where some behaviors were organized by early hormone exposure and others were not. Our results confirm that in birds, chromosomal sex plays a major–and perhaps dominant–role in shaping gene expression patterns during neurodevelopment via cell-autonomous sex identity. This is in contrast to what is seen in mammals, where early hormone exposure shapes sexual differentiation of the brain in consistent ways, though this is more a matter of degree than kind. Sex chromosome effects certainly occur in mammals, as well, and this phenomenon is an area of active research ^23,24,74^. In chickens, the Z chromosome accounts for approximately 7% of the entire genome and contains more genes than the mammalian X chromosome^75^. So it is unsurprising that the lack of gene dosage compensation leads to substantial sex chromosome dependent gene expression.

Our approach here allows us to partially disentangle the separate contributions of sex chromosomes and hormones, which would normally be reinforcing in untreated animals. Sexual differentiation is a canalizing process, where the developmental stability of sex-related phenotypes is expected to be adaptive ^76,77^. Our results clearly suggest that sexually differentiated phenotypes in birds can be shaped by sex chromosome complement alone, organizational effects of hormones alone, but more often by both together. However, these developmental processes are complex and likely play out differently by tissue and even cell-type. This also explains the remarkable sex diversity observed in birds, including observations of naturally-occurring “crowing hens” ^78^ and gynandromorphs ^29,31,32^. Thus, our results provide insight into the dynamic, complex process by which sexual differentiation occurs in the brains of birds, illuminating the true sex diversity in the natural world.

## Resource availability

RNA sequencing data have been deposited in NCBI BioProject under accession code PRJNA1533991.

## Acknowledgements

We would like to acknowledge the North Carolina State University Animal & Poultry Teaching Unit for providing fertilized chicken eggs. We are also extremely grateful for lab managers Jessica Spillman and Georgiana Walters, as well as members of the Baran lab, for their assistance with egg care. VM was supported for two summers of research by the Davidson College Martin ’85 and Lisa ’86 Foil Neuroscience Research Award.

## STAR Methods

### Chicken eggs

We used a total of 63 fertilized White Leghorn chicken (*Gallus gallus domesticus*) eggs from North Carolina State University Animal & Poultry Teaching Unit (Raleigh, NC). We split the 63 eggs into three batches and began incubating one batch per day over the span of three days. Eggs were incubated at 37.8°C under humid conditions and rotated once every hour using a 1502 Sportsman Incubator (Savannah, GA). All procedures were approved by the Davidson College Institutional Animal Care and Use Committee (Protocol 15-24-02).

### Experimental and control treatments

We injected eggs on E4 (the fourth day after incubation had begun, roughly Hamburger Hamilton Stage 23)^79^ with either an experimental or control treatment. After poking a small hole in the shell of the eggs using a teasing needle, we injected the experimental group (n = 27) with 1mg of fadrozole (F3806 from Sigma-Aldrich, Saint-Louis, MO) dissolved in 0.1 mL of sterile physiological saline solution (10mg/mL concentration). The control group (n = 36) received 0.1 mL of sterile physiological saline solution. Following injection, eggs were sealed with tape and returned to the incubator.

### Dissection, tissue collection & gonadal sexing

On E11 of egg incubation (Hamburger Hamilton Stage 37)^79^, we dissected the eggs and humanely euthanized embryos via rapid decapitation. After decapitation, we flash-froze whole heads on dry ice and stored these samples at −80C. We collected blood immediately following decapitation at the site of the freshly severed carotid artery using capillary tubes. We then centrifuged these samples, working over ice, and pipetted the separated plasma into microcentrifuge tubes, which were stored at −20°C. We collected additional blood samples (or tissue samples in the case of stunted embryonic development) of ∼20μl in microcentrifuge tubes pre-filled with 180μL of ATL buffer from a Qiagen DNeasy Blood & Tissue kit (Cat. No. 69504). These samples were stored at 5°C for later use in chromosomal sexing. Finally, we dissected embryos and imaged and phenotyped their gonads visually using a dissecting microscope (Figure 1C). Testes were defined as gonads which appeared symmetrical, long, and slender, while the ovary at this stage is large and bulbous on the anatomical left side with a degrading undifferentiated gonad on the anatomical right. Gonads were designated as ovotestes if the gonads were in between testes or ovaries, often slightly asymmetrical with a rounder and bigger right gonad than typical testes, but smaller and more slender than a typical ovary ^80^. Gonadal phenotypes were later confirmed blindly using anonymized images.

### Chromosomal sexing

Using the previously collected blood and tissue samples, we performed DNA extraction using a Qiagen DNeasy Blood & Tissue kit (Cat. No. 69504) following manufacturer’s instructions. We then carried out a PCR using the CHD1 primers as described in ^81^ : Fw (2550F): GTTACTGATTCGTCTACGAGA, Rv (2718R): ATTGAAATGATCCAGTGCTTG. Each 20μl PCR reaction contained 7μl RNAse-free H_2_0; 1μl 5μM F+R primer mix; 10μl DreamTaq Hot Start Green PCR Master Mix (Cat. No. K9021); and 2μl of sample DNA. We used the following thermocycler conditions: an initial 5min denaturation step at 95°C; followed by a “touch-down” scheme of denaturation at 94°C for 30 sec, annealing for 30 sec where the temperature decreased 1°C per cycle from 60°C until it reached a temperature of 51°C (10 cycles), and extension 72°C for 1min; 30 cycles where each consisted of denaturation at 94°C for 30 sec, annealing at 50°C for 30 sec, and extension at 72°C for 1 min; followed by a final extension step at 72°C for 5 min. We analyzed PCR products by 1.25% agarose gel electrophoresis (in 1% in TBE buffer) and visualized them using SYBR Safe DNA Gel Stain (Invitrogen, Cat. No. S33102).

### Steroid analysis

Plasma steroids were measured using an ultrasensitive and specific LC-MS/MS assay. The steroid panel included pregnenolone, progesterone, dehydroepiandrosterone, androstenedione, androstenediol, testosterone, 5α-dihydrotestosterone, 17β-estradiol, estriol, corticosterone, 11-deoxycortisol, cortisol, and cortisone.

Steroids were extracted from plasma samples (20 μL) using liquid-liquid extraction, followed by derivatization with 1,2-dimethylimidazole-5-sulfonyl-chloride (DMIS) to increase sensitivity for estrogens ^82–84^. Each sample was placed along with five zirconium ceramic oxide beads (1.4 mm diameter) in a 2 mL polypropylene tube, and then 50 μL of deuterated internal standards (pregnenolone-d4, progesterone-d9, dehydroepiandrosterone-d6, testosterone-d5, 17β-estradiol-d4, corticosterone-d8, cortisol-d4; C/D/N Isotopes Inc., Pointe-Claire, QC, Canada) were added to track recovery and adjust for matrix interference. Samples received 1 mL of high-performance liquid chromatography (HPLC)-grade acetonitrile, were vortexed, and then homogenized at 4 m/s for 30 s using a bead mill homogenizer. After centrifugation at 16,100 g for 15 min, 1 mL of supernatant was collected and transferred to a pre-cleaned 12 × 75 mm borosilicate glass culture tube. Next, samples received 500 μL of HPLC-grade hexane, were vortexed, and then centrifuged at 3200 g for 2 min. Hexane was removed and discarded, and acetonitrile was evaporated at 60°C for 45 min in a vacuum centrifuge. The dried samples were placed in a wet ice bath and resuspended in 30 μL of sodium bicarbonate buffer (50 mM, pH 10.5) and then 20 μL of 1 mg/mL DMIS in HPLC-grade acetone was added. Samples were vortexed, centrifuged at 3200 g for 1 min, and transferred to glass inserts in LC vials. Vials were capped and incubated at 60°C for 15 min, cooled at 4°C for 15 min, centrifuged at 3200 g for 1 min, and stored at −20°C until injection for LC-MS/MS analysis.

Samples were processed alongside a calibration curve of known amounts of steroid standards made from certified reference standards (Cerilliant, TX, USA) or prepared in-house from commercial steroids (Steraloids, RI, USA) in 50% HPLC-grade methanol in MilliQ water. Lower limits of quantification (LLOQs) were 0.01 pg/tube for estriol, 0.03 pg/tube for 17β-estradiol and estrone, 0.04 pg/tube for testosterone, 0.1 pg/tube for androstenedione, 0.2 pg/tube for progesterone and corticosterone, 0.4 pg/tube for 5α-dihydrotestosterone and cortisol, 0.8 pg/tube for dehydroepiandrosterone and 11-deoxycortisol, 5 pg/tube for pregnenolone and androstenediol, and 20 pg/tube for cortisone. All analytes demonstrated a linear range up to 1000 pg/tube. Of these analytes, only 17β-estradiol, estrone, testosterone, androstenedione, progesterone, corticosterone, and 11-deoxycortisol were above the LLOQs in plasma samples, whereas other steroids were below the LLOQs.

Assay accuracy and precision were evaluated by measuring triplicates of blanks (internal standards only), double blanks (without standards or internal standards), and quality controls (0.2 pg, 2 pg, and 20 pg) prepared in neat solution. All blanks and double blanks had non-detectable analyte levels, and quality controls were acceptable (within 20% of target values).

To assess derivatization efficiency, underivatized standards were included to quantify underivatized estrogens. Underivatized standards received 20 μL of acetone without DMIS. The derivatization efficiency was greater than 99%.

Steroids were quantified using a QTRAP 6500 UHPLC-MS/MS system (Sciex LLC, Framingham, MA, USA) ^82,84^. Samples were loaded into a refrigerated autosampler at 15 °C, and 35 μL of each sample was injected into a Nexera X2 UHPLC system (Shimadzu Corp., Kyoto, Japan) and passed through a KrudKatcher ULTRA HPLC In-Line Filter (Phenomenex, Torrance, CA, USA). Analytes were then separated on a 2.1 × 50 mm Kinetex® 2.6 μM EVO C18 100 A° LC column at 40°C using a binary gradient mobile system with 0.1 mM ammonium fluoride in MilliQ water as mobile phase A and HPLC-grade methanol as mobile phase B (MPB) set at a 0.4 mL/min flow rate. Total run time was 14.9 min during which MPB was 10% from 0 to 0.5 min and increased to 42% at 0.5 min, 60% at 4.0 min, 70% at 9.4 min, and 98% at 9.5 min until 11.9 min. A column wash was then carried out at 98% MPB until 13.4 min, and the MPB was then returned to 10% until the end of the run. Before and after each injection, the autoinjector needle was rinsed externally with 100% HPLC-grade methanol. Two scheduled multiple reaction monitoring transitions were used for each analyte, and one transition was used for each internal standard. All analytes were acquired in positive electrospray ionization mode, except for underivatized estrogens, which were acquired in negative mode.

### Brain tissue sample collection & processing

We mounted the whole frozen heads collected during dissection using a cryostat. Samples were mounted to the chuck at a ∼305° angle and sliced at a ∼105° angle coronally at 60μm until hypothalamus tissue was exposed, as indicated by the embryonic chicken brain atlas (Chick Embryo Atlas E10.5 Frontal) ^85^. Once sufficient hypothalamus was exposed we used a 1mm tissue punch at a ∼60° angle to collect a tissue sample which penetrated the whole of the hypothalamus (Supplementary Figure 1). We then collected samples in microcentrifuge tubes filled with 20μl of Zymo Research DNA/RNA Shield tissue reagent (Zymo Research, #R1100). Following centrifugation, we stored samples at −20°C for use in RNA sequencing.

### RNA sequencing

Of the tissue samples collected from the whole frozen heads, we selected 20 samples in which we had the highest confidence of tissue punch placement and plasma sample quality for RNA sequencing. We performed RNA extraction using a Qiagen AllPrep DNA/RNA Mini Kit (Cat. No. 80204). We used the protocol “Simultaneous Purification of Genomic DNA and Total RNA from Animal Tissues” following guidelines for RNAprotect-stabilized tissues.

Once RNA extraction was completed, we stored samples at −80°C and shipped them on dry ice to Novogene Corporation, Inc. (Sacramento, CA) for quality control screening, library preparation, and RNA sequencing. The concentration and integrity of all RNA samples was determined using an Agilent 5400 Bioanalyzer system and all samples proved to be of high quality (mean RNA concentration 43.3 ±12.0 ng/µl; RIN = 9.77 ± 0.06). Next, mRNA was purified from total RNA using poly-T oligo attached magnetic beads. After cDNA synthesis, end repair, A-tailing, adaptor ligation, size selection, amplification, and purification using the Vazyme VAHTS Universal V10 RNA-seq Library Prep Kit for Illumina (Cat no. NR606), the cDNA libraries were pooled in equimolar concentrations and were sequenced on an Illumina Novaseq X Plus instrument (Illumina Inc., CA, USA). We sequenced 150bp paired-end reads with a depth of 6.4 ± 0.93. Raw sequence reads are available via NCBI at BioProject ID PRJNA1533991.

### Bioinformatic pipeline & processing

With the raw sequencing data from Novogene, we assessed the quality of the raw reads using *FastQC* v0.12. ^86^. Once we had confirmed sufficient quality of the reads, we used *HISAT2* v2.2.1 ^87^ to align reads to the GRCg6a version of the chicken genome downloaded from the Ensembl Site. Following alignment with the genome we used *Qualimaps* v2.3 ^88^ to confirm the quality of the overall alignment of our reads to the reference genome. We then used *StringTie* v3.0.3 ^89^ to quantify the expression of individual genes.

### Gene expression analysis

Working in R Statistical Software v4.2.1, we used *DESeq2* ^90^ with the default normalization settings to identify genes exhibiting differential expression by treatment, sex chromosome, and an interaction of the two. We used the following DESeq2 model: ∼ Chromosome + Treatment + Chromosome:Treatment. The chromosomal factor was leveled ZZ to ZW and the treatment factor was leveled saline to fadrozole. The five resulting contrasts of this model compared: ZZ to ZW control individuals, ZZ to ZW fadrozole-treated individuals, ZZ control to ZZ fadrozole-treated individuals, ZW control to ZW fadrozole-treated individuals, and finally a contrast for the interaction term (Figure 1B). The two contrasts comparing chromosomal sexes allow us to find the main effects of chromosomal sex within both treatment groups, while the two comparing treatment groups allow us to find the main effects of treatment within both chromosomal sexes. The interaction term captures genes in which the effect of treatment differs between and depends on chromosomal sex. We elected to use an unadjusted p-value of < 0.01. While less stringent and potentially more prone to error, this approach aligns with the exploratory nature of the current study ^91^. Using this significance threshold, lists of differentially expressed genes were created for each contrast with respective fold-difference and significance values for each gene (Supplementary Table 2).

### Weighted-gene coexpression network analysis

Given that we have an adequate sample size (n = 20), we performed weighted gene coexpression network analysis (WGCNA) using the *WGCNA* R package ^92^. For this analysis we used a pairwise comparison model producing six contrasts for comparison between each of the four sample groups. After inspecting sample clustering to confirm there were no outlier samples, we ensured data quality by filtering genes with missing values or low variance using a variance threshold of 0.4. Soft thresholding power (*β*) determines how strongly high versus low correlations between genes are weighted. We determined *β* should be set to 12 based on a scale-free topology with R^2^ = 0.85 and low mean connectivity near 0. We then used the blockwiseModules function to calculate gene pair correlations and adjacencies, compute the Topological Overlap Matrix (TOM), perform hierarchical clustering, and detect modules using dynamic tree cutting. Within this function we used the predetermined *β, a* minimum module size of 30, a merge cut height of 0.25, and signed network type to preserve direction of correlation.

Using these parameters, 24 distinct modules with highly correlated expression patterns were identified from a set of 14,706 filtered genes, and a 25th module (grey) held unassigned genes. We then calculated the module eigengene (ME) for each module, which is the first principal component of gene expression levels for all genes in a module. MEs represent the overall expression pattern of genes within a module. With the MEs calculated we performed an analysis to find the mean values of each module’s ME by sample group and plotted the resulting distribution and standard error, which was calculated based on individual sample ME values (Figure 3B). Also using the ME data we calculated module-trait associations (MTA), mapping the strength and direction of gene expression correlations between each module and each contrast relating to ME expression, and determining whether there is a significant (*p* < 0.05) correlation between them (Figure 3A). While the correlations between each module and the SZZ vs FZW and FZZ vs SZW contrasts were calculated, these correlations were not presented because they provide information more difficult to interpret biological meaning from compared to other contrasts. We then created a module membership (MM) matrix by calculating the Pearson correlation between gene expression levels and each module eigengene. The closer an MM is to 1, the higher the degree of connections between a specific gene and genes in other modules. The gene significance (GS) was also calculated for each module in relation to each contrast to measure how strongly a gene’s expression correlates with each trait. A GS > 0.05 indicates a gene is individually associated with a trait. Correlations between MM and GS were calculated for each module and each contrast to find combinations with strong positive correlation. We then identified hub genes within each module, which are those with high MM and GS, and strong correlations to other genes within the module using a threshold of 0.05 to filter adjacencies within the TOM. Using the top 20 hub genes by adjacencies for each module we created a co-expression network plot to show the relationships between modules, including how they cluster together (Figure 3C) Finally, we quantified and plotted the number of genes within each module according to location on the Z, W, or autosomal chromosomes to identify whether there was over-or underrepresentation of chromosomes within each module (Figure 3D).

### Gene Ontology analysis

To decipher the potential underlying biological mechanisms creating the patterns of gene expression seen in our samples, we performed gene ontology (GO) analysis using *ShinyGO* v0.86.0 (Ge et al., 2020). We identified GO terms, which are biological categories statistically overrepresented by gene products within a list of candidate genes, for both differential expression and WGCNA data. For the differential expression data, GO analysis was performed on gene lists for each of the contrasts’ up- and down-regulated genes. For WGCNA data, GO analysis was performed on the list of genes contained within each module. For GO analysis on the WGCNA dataset, a background of the 14,706 filtered genes was used, while for the differential expression data, no background was used because this dataset pulls from the full chicken genome. For both sets of gene lists, we identified GO terms classified under biological processes, cellular component, molecular functions, Kyoto Encyclopedia of Genes & Genomics (KEGG), Protein Analysis Through Evolutionary Relationships (PANTHER), Reactome, and Wikipathways. Each GO term identified was assigned a False Discovery Rate value and a fold-enrichment value indicating the scale of the overrepresentations of genes for the GO term in the dataset.

### ChIP-Seq Comparative Analysis

To better understand the direct impacts of steroid hormones on gene expression within our differentially expressed gene lists and WGCNA modules, we compared these sets of genes with those previously identified by past ChIP-Seq studies of AR, ERɑ, ERꞵ, MR, and GR ^57–60^. We used a Chi-Square Test of Independence for each gene list set to determine if there was a statistically significant association between the gene list and the ChIP-Seq gene list. Using a Bonferroni correction, we only considered associations with a p-value of less than 0.0014 to be significant, given 35 total comparisons. In addition, we considered which modules from the WGCNA contributed >10% of the overall *χ*^2^ value as those with under-or overrepresentation.

## References

1. McCarthy, M.M., and Arnold, A.P. (2011). Reframing sexual differentiation of the brain. Nat. Neurosci. 14, 677–683. 10.1038/nn.2834.

2. Morrish, B.C., and Sinclair, A.H. (2002). Vertebrate sex determination: many means to an end. Reproduction 124, 447–457. 10.1530/rep.0.1240447.

3. Massa, M.G., Aghi, K., and Hill, M. (2023). Deconstructing sex: Strategies for undoing binary thinking in neuroendocrinology and behavior. Horm. Behav. 156, 105441. 10.1016/j.yhbeh.2023.105441.

4. Smiley, K.O., Munley, K.M., Aghi, K., Lipshutz, S.E., Patton, T.M., Pradhan, D.S., Solomon-Lane, T.K., and Sun, S.D. (2024). Sex diversity in the 21st century: Concepts, frameworks, and approaches for the future of neuroendocrinology. Horm. Behav. 157, 105445. 10.1016/j.yhbeh.2023.105445.

5. Wilhelm, D., Palmer, S., and Koopman, P. (2007). Sex determination and gonadal development in mammals. Physiol. Rev. 87, 1–28. 10.1152/physrev.00009.2006.

6. Koopman, P., Gubbay, J., Vivian, N., Goodfellow, P., and Lovell-Badge, R. (1991). Male development of chromosomally female mice transgenic for Sry. Nature 351, 117–121. 10.1038/351117a0.

7. McCarthy, M.M. (2008). Estradiol and the developing brain. Physiol. Rev. 88, 91–134. 10.1152/physrev.00010.2007.

8. Balthazart, J., and Ball, G. (2013). Brain aromatase, estrogens, and behavior (OUP USA).

9. Ioannidis, J., Taylor, G., Zhao, D., Liu, L., Idoko-Akoh, A., Gong, D., Lovell-Badge, R., Guioli, S., McGrew, M.J., and Clinton, M. (2021). Primary sex determination in birds depends on DMRT1 dosage, but gonadal sex does not determine adult secondary sex characteristics. Proc. Natl. Acad. Sci. U. S. A.. 118, e2020909118. 10.1073/pnas.2020909118.

10. Hirst, C.E., Major, A.T., Ayers, K.L., Brown, R.J., Mariette, M., Sackton, T.B., and Smith, C.A. (2017). Sex reversal and comparative data undermine the W chromosome and support Z-linked DMRT1 as the regulator of gonadal sex differentiation in birds. Endocrinology 158, 2970–2987. 10.1210/en.2017-00316.

11. Chue, J., and Smith, C.A. (2011). Sex determination and sexual differentiation in the avian model. FEBS J. 278, 1027–1034. 10.1111/j.1742-4658.2011.08032.x.

12. Ayers, K.L., Smith, C.A., and Lambeth, L.S. (2013). The molecular genetics of avian sex determination and its manipulation. GENESIS 51, 325–336. 10.1002/dvg.22382.

13. Smith, C.A., Roeszler, K.N., Ohnesorg, T., Cummins, D.M., Farlie, P.G., Doran, T.J., and Sinclair, A.H. (2009). The avian Z-linked gene DMRT1 is required for male sex determination in the chicken. Nature 461, 267–271. 10.1038/nature08298.

14. Andrews, J.E., Smith, C.A., and Sinclair, A.H. (1997). Sites of estrogen receptor and aromatase expression in the chicken embryo. Gen. Comp. Endocrinol. 108, 182–190. 10.1006/gcen.1997.6978.

15. Nomura, O., Nakabayashi, O., Nishimori, K., Yasue, H., and Mizuno, S. (1999). Expression of five steroidogenic genes including aromatase gene at early developmental stages of chicken male and female embryos. J. Steroid Biochem. Mol. Biol. 71, 103–109. 10.1016/S0960-0760(99)00127-2.

16. Smith, C.A., Katz, M., and Sinclair, A.H. (2003). DMRT1 is upregulated in the gonads during female-to-male sex reversal in ZW chicken embryos. Biol. Reprod. 68, 560–570. 10.1095/biolreprod.102.007294.

17. Scheib, D. (1983). Effects and role of estrogens in avian gonadal differentiation. In Mechanisms of Gonadal Differentiation in Vertebrates: Contributions of an EMBO-Workshop held in Freiburg, November 5–8, 1982, U. Müller and W. W. Franke, eds. (Springer), pp. 87–92. 10.1007/978-3-642-69150-8_15.

18. Itoh, Y., Melamed, E., Yang, X., Kampf, K., Wang, S., Yehya, N., Van Nas, A., Replogle, K., Band, M.R., Clayton, D.F., et al. (2007). Dosage compensation is less effective in birds than in mammals. J. Biol. 6, 2. 10.1186/jbiol53.

19. Ellegren, H., Hultin-Rosenberg, L., Brunström, B., Dencker, L., Kultima, K., and Scholz, B. (2007). Faced with inequality: chicken do not have a general dosage compensation of sex-linked genes. BMC Biol. 5, 40. 10.1186/1741-7007-5-40.

20. Arnold, A.P., Itoh, Y., and Melamed, E. (2008). A bird’s-eye view of sex chromosome dosage compensation. Annu. Rev. Genom. Hum. Genet. 9, 109–127. 10.1146/annurev.genom.9.081307.164220.

21. Wolf, J.B., and Bryk, J. (2011). General lack of global dosage compensation in ZZ/ZW systems? Broadening the perspective with RNA-seq. BMC Genomics 12, 91. 10.1186/1471-2164-12-91.

22. Sun, D., Layman, T.S., Jeong, H., Chatterjee, P., Grogan, K., Merritt, J.R., Maney, D.L., and Yi, S.V. (2021). Genome-wide variation in DNA methylation linked to developmental stage and chromosomal suppression of recombination in white-throated sparrows. Mol. Ecol. 30, 3453–3467. 10.1111/mec.15793.

23. Arnold, A.P. (2009). Mouse models for evaluating sex chromosome effects that cause sex differences in non-gonadal tissues. J. Neuroendocrinol. 21, 377–386. 10.1111/j.1365-2826.2009.01831.x.

24. Arnold, A.P., and Chen, X. (2009). What does the “four core genotypes” mouse model tell us about sex differences in the brain and other tissues? Front. Neuroendocrinol. 30, 1–9. 10.1016/j.yfrne.2008.11.001.

25. Clinton, M., Zhao, D., Nandi, S., and McBride, D. (2012). Evidence for avian cell autonomous sex identity (CASI) and implications for the sex-determination process? Chromosome Res. 20, 177–190. 10.1007/s10577-011-9257-9.

26. Major, A.T., and Smith, C.A. (2016). Sex reversal in birds. Sex. Dev. 10, 288–300. 10.1159/000448365.

27. Zhang, X., Li, J., Chen, S., Yang, N., and Zheng, J. (2023). Overview of avian sex reversal. Int. J. Mol. Sci. 24. 10.3390/ijms24098284.

28. Brunström, B., Axelsson, J., Mattsson, A., and Halldin, K. (2009). Effects of estrogens on sex differentiation in Japanese quail and chicken. Gen. Comp. Endocrinol. 163, 97–103. 10.1016/j.ygcen.2009.01.006.

29. Agate, R.J., Grisham, W., Wade, J., Mann, S., Wingfield, J., Schanen, C., Palotie, A., and Arnold, A.P. (2003). Neural, not gonadal, origin of brain sex differences in a gynandromorphic finch. Proc. Natl. Acad. Sci. U. S. A.. 100, 4873–4878. 10.1073/pnas.0636925100.

30. Zhao, D., McBride, D., Nandi, S., McQueen, H.A., McGrew, M.J., Hocking, P.M., Lewis, P.D., Sang, H.M., and Clinton, M. (2010). Somatic sex identity is cell autonomous in the chicken. Nature 464, 237–242. 10.1038/nature08852.

31. Morris, K.R., Hirst, C.E., Major, A.T., Ezaz, T., Ford, M., Bibby, S., Doran, T.J., and Smith, C.A. (2018). Gonadal and endocrine analysis of a gynandromorphic chicken. Endocrinology 159, 3492–3502. 10.1210/en.2018-00553.

32. Cao, Y., Huang, Z., Hu, W., Zhang, Y., Zhang, K., Zuo, J., Zou, Y., and Meng, W. (2026). Cell-autonomous determination of sex dimorphism in neuromodulation of a gynandromorphic Zebra Finch. Avian Res. 17, 100327. 10.1016/j.avrs.2025.100327.

33. Lee, H.J., Seo, M., Choi, H.J., Rengaraj, D., Jung, K.M., Park, J.S., Lee, K.Y., Kim, Y.M., Park, K.J., Han, S.T., et al. (2021). DMRT1 gene disruption alone induces incomplete gonad feminization in chicken. FASEB J. 35, e21876. 10.1096/fj.202100902R.

34. Adkins, E.K. (1975). Hormonal basis of sexual differentiation in the Japanese quail. J. Comp. Physiol. Psychol. 89, 61–71. 10.1037/h0076406.

35. Balthazart, J., and Ball, G.F. (1995). Sexual differentiation of brain and behavior in birds. Trends Endocrinol. Metab. 6, 21–29. 10.1016/1043-2760(94)00098-O.

36. Sayag, N., Snapir, N., Robinzon, B., Arnon, E., Halawani, M.E.E., and Grimm, V.E. (1989). Embryonic sex steroids affect mating behavior and plasma LH in adult chickens. Physiol. Behav. 45, 1107–1112. 10.1016/0031-9384(89)90095-4.

37. Panzica, G.C., Viglietti-Panzica, C., and Balthazart, J. (1996). The sexually dimorphic medial preoptic nucleus of quail: A key brain area mediating steroid action on male sexual behavior. Front. Neuroendocrinol. 17, 51–125. 10.1006/frne.1996.0002.

38. Panzica, G.C., Castagna, C., Viglietti-Panzica, C., Russo, C., Tlemçani, O., and Balthazart, J. (1998). Organizational effects of estrogens on brain vasotocin and sexual behavior in quail. J. Neurobiol. 37, 684–699. 10.1002/(SICI)1097-4695(199812)37:4%3C684::AID-NEU15%3E3.0.CO;2-U.

39. Suzuki, M., and Tsukahara, S. (2025). *In ovo* exposure to ethynylestradiol demasculinizes the vasotocin system in the medial preoptic nucleus in male Japanese quail. Neurosci. Lett. 864, 138310. 10.1016/j.neulet.2025.138310.

40. Cornil, C.A., Ball, G.F., Balthazart, J., and Charlier, T.D. (2011). Organizing effects of sex steroids on brain aromatase activity in quail. PLOS ONE 6, e19196. 10.1371/journal.pone.0019196.

41. Panzica, G.C., Viglietti-Panzica, C., Calagni, M., Anselmetti, G.C., Schumacher, M., and Balthazart, J. (1987). Sexual differentiation and hormonal control of the sexually dimorphic medial preoptic nucleus in the quail. Brain. Res. 416, 59–68. 10.1016/0006-8993(87)91496-X.

42. Delage, C.I., and Cornil, C.A. (2020). Estrogen-dependent sex difference in microglia in the developing brain of Japanese quail (*Coturnix japonica*). Dev. Neurobiol. 80, 239–262. 10.1002/dneu.22781.

43. Cornil, C.A., and Balthazart, J. (2023). Contribution of birds to the study of sexual differentiation of brain and behavior. Horm. Behav. 155, 105410. 10.1016/j.yhbeh.2023.105410.

44. Gurney, M.E., and Konishi, M. (1980). Hormone-induced sexual differentiation of brain and behavior in zebra finches. Science 208, 1380–1383. 10.1126/science.208.4450.1380.

45. Pohl-Apel, G., and Sossinka, R. (1984). Hormonal determination of song capacity in females of the zebra finch: Critical phase of treatment. Z. Tierpsychol. 64, 330–336. 10.1111/j.1439-0310.1984.tb00367.x.

46. Simpson, H.B., and Vicario, D.S. (1991). Early estrogen treatment of female zebra finches masculinizes the brain pathway for learned vocalizations. J. Neurobiol. 22, 777–793. 10.1002/neu.480220711.

47. Simpson, H.B., and Vicario, D.S. (1991). Early estrogen treatment alone causes female zebra finches to produce learned, male-like vocalizations. J. Neurobiol. 22, 755–776. 10.1002/neu.480220710.

48. Adkins-Regan, E., and Ascenzi, M. (1990). Sexual differentiation of behavior in the zebra finch: Effect of early gonadectomy or androgen treatment. Horm. Behav. 24, 114–127. 10.1016/0018-506X(90)90031-R.

49. Adkins-Regan, E., Mansukhani, V., Seiwert, C., and Thompson, R. (1994). Sexual differentiation of brain and behavior in the zebra finch: Critical periods for effects of early estrogen treatment. J. Neurobiol. 25, 865–877. 10.1002/neu.480250710.

50. Schlinger, B.A., and Arnold, A.P. (1992). Plasma sex steroids and tissue aromatization in hatchling zebra finches: implications for the sexual differentiation of singing behavior. Endocrinology 130, 289–299. 10.1210/en.130.1.289.

51. Hutchison, J.B., Wingfield, J.C., and Hutchison, R.E. (1984). Sex differences in plasma concentrations of steroids during the sensitive period for brain differentiation in the zebra finch. J. Endocrinol. 103, 363–369. 10.1677/joe.0.1030363.

52. Adkins-Regan, E., Abdelnabi, M., Mobarak, M., and Ottinger, M.A. (1990). Sex steroid levels in developing and adult male and female zebra finches (*Poephila guttata*). Gen. Comp. Endocrinol. 78, 93–109. 10.1016/0016-6480(90)90051-M.

53. Choe, H.N., Tewari, J., Zhu, K.W., Davenport, M., Matsunami, H., and Jarvis, E.D. (2021). Estrogen and sex-dependent loss of the vocal learning system in female zebra finches. Horm. Behav. 129, 104911. 10.1016/j.yhbeh.2020.104911.

54. Davenport, M., Choe, H.N., Matsunami, H., and Jarvis, E. (2025). Sex chromosome gene expression associated with vocal learning following hormonal manipulation in female zebra finches. eLife 12, RP89425. 10.7554/eLife.89425.

55. Elbrecht, A., and Smith, R.G. (1992). Aromatase enzyme activity and sex determination in chickens. Science 255, 467–470. 10.1126/science.1734525.

56. Wobbrock, J.O., Findlater, L., Gergle, D., and Higgins, J.J. (2011). The aligned rank transform for nonparametric factorial analyses using only anova procedures. In Proceedings of the SIGCHI Conference on Human Factors in Computing Systems CHI ’11. (Association for Computing Machinery), pp. 143–146. 10.1145/1978942.1978963.

57. Quartier, A., Chatrousse, L., Redin, C., Keime, C., Haumesser, N., Maglott-Roth, A., Brino, L., Gras, S.L., Benchoua, A., Mandel, J.-L., et al. (2018). Genes and pathways regulated by androgens in human neural cells, potential candidates for the male excess in autism spectrum disorder. Biol. Psychiatry 84, 239–252. 10.1016/j.biopsych.2018.01.002.

58. Gegenhuber, B., Wu, M.V., Bronstein, R., and Tollkuhn, J. (2022). Gene regulation by gonadal hormone receptors underlies brain sex differences. Nature 606, 153–159. 10.1038/s41586-022-04686-1.

59. Grober, O.M., Mutarelli, M., Giurato, G., Ravo, M., Cicatiello, L., De Filippo, M.R., Ferraro, L., Nassa, G., Papa, M.F., Paris, O., et al. (2011). Global analysis of estrogen receptor beta binding to breast cancer cell genome reveals an extensive interplay with estrogen receptor alpha for target gene regulation. BMC Genomics 12, 36. 10.1186/1471-2164-12-36.

60. Mifsud, K.R., Kennedy, C.L.M., Salatino, S., Sharma, E., Price, E.M., Haque, S.N., Gialeli, A., Goss, H.M., Panchenko, P.E., Broxholme, J., et al. (2021). Distinct regulation of hippocampal neuroplasticity and ciliary genes by corticosteroid receptors. Nat. Commun. 12, 4737. 10.1038/s41467-021-24967-z.

61. Simkins, J.W., Joseph, A.E., Bonier, F., and Benowitz-Fredericks, Z.M. (2020). Prenatal aromatase inhibition alters postnatal immunity in domestic chickens (*Gallus gallus*). Gen. Comp. Endocrinol. 294, 113497. 10.1016/j.ygcen.2020.113497.

62. Woods, J.E., and Brazzill, D.M. (1981). Plasma 17β-estradiol levels in the chick embryo. Gen. Comp. Endocrinol. 44, 37–43. 10.1016/0016-6480(81)90353-1.

63. Tanabe, Y., Saito, N., and Nakamura, T. (1986). Ontogenetic steroidogenesis by testes, ovary, and adrenals of embryonic and postembryonic chickens (*Gallus domesticus*). Gen. Comp. Endocrinol. 63, 456–463. 10.1016/0016-6480(86)90146-2.

64. Strushkevich, N., Gilep, A.A., Shen, L., Arrowsmith, C.H., Edwards, A.M., Usanov, S.A., and Park, H.-W. (2013). Structural insights into aldosterone synthase substrate specificity and targeted inhibition. Mol. Endocrinol. 27, 315–324. 10.1210/me.2012-1287.

65. Brixius-Anderko, S., and Scott, E.E. (2019). Structure of human cortisol-producing cytochrome P450 11B1 bound to the breast cancer drug fadrozole provides insights for drug design. J. Biol. Chem. 294, 453–460. 10.1074/jbc.RA118.006214.

66. Bordt, E.A., Ceasrine, A.M., and Bilbo, S.D. (2020). Microglia and sexual differentiation of the developing brain: A focus on ontogeny and intrinsic factors. Glia 68, 1085–1099. 10.1002/glia.23753.

67. VanRyzin, J.W., Marquardt, A.E., Pickett, L.A., and McCarthy, M.M. (2020). Microglia and sexual differentiation of the developing brain: A focus on extrinsic factors. Glia 68, 1100–1113. 10.1002/glia.23740.

68. Jensen, C.J., Massie, A., and De Keyser, J. (2013). Immune players in the CNS: The astrocyte. J. Neuroimmune. Pharmacol. 8, 824–839. 10.1007/s11481-013-9480-6.

69. Saldanha, C.J. (2021). Glial estradiol synthesis after brain injury. Curr. Opin. Endocr. Metab. Res. 21, 100298. 10.1016/j.coemr.2021.100298.

70. Ma, Y., Guo, H., Zhang, L., Tao, L., Yin, A., Liu, Z., Li, Y., Dong, H., Xiong, L., and Hou, W. (2016). Estrogen replacement therapy-induced neuroprotection against brain ischemia-reperfusion injury involves the activation of astrocytes via estrogen receptor β. Sci. Rep. 6, 21467. 10.1038/srep21467.

71. Itoh, N., Itoh, Y., Meyer, C.E., Suen, T.T., Cortez-Delgado, D., Rivera Lomeli, M., Wendin, S., Somepalli, S.S., Golden, L.C., MacKenzie-Graham, A., et al. (2023). Estrogen receptor beta in astrocytes modulates cognitive function in mid-age female mice. Nat. Commun. 14, 6044. 10.1038/s41467-023-41723-7.

72. Klinge, C.M. (2020). Estrogenic control of mitochondrial function. Redox Biol. 31, 101435. 10.1016/j.redox.2020.101435.

73. Lassiter, K., Dridi, S., Greene, E., Kong, B., and Bottje, W.G. (2018). Identification of mitochondrial hormone receptors in avian muscle cells. Poult. Sci. 97, 2926–2933. 10.3382/ps/pey126.

74. Snell, D.M., and Turner, J.M.A. (2018). Sex chromosome effects on male–female differences in mammals. Curr Biol 28, R1313–R1324. 10.1016/j.cub.2018.09.018.

75. Bellott, D.W., Skaletsky, H., Pyntikova, T., Mardis, E.R., Graves, T., Kremitzki, C., Brown, L.G., Rozen, S., Warren, W.C., Wilson, R.K., et al. (2010). Convergent evolution of chicken Z and human X chromosomes by expansion and gene acquisition. Nature 466, 612–616. 10.1038/nature09172.

76. Waddington, C.H. (1942). Canalization of development and the inheritance of acquired characters. Nature 150, 563–565. 10.1038/150563a0.

77. Siegal, M.L., and Bergman, A. (2002). Waddington’s canalization revisited: Developmental stability and evolution. Proc. Natl. Acad. Sci. U. S. A.. 99, 10528–10532. 10.1073/pnas.102303999.

78. Forbes, T.R. (1947). The crowing hen: Early observations on spontaneous sex reversal in birds. Yale J. Biol. Med. 19, 955–970.

79. Hamburger, V., and Hamilton, H.L. (1951). A series of normal stages in the development of the chick embryo. J. Morphol. 88, 49–92. 10.1002/jmor.1050880104.

80. Lambeth, L.S., Cummins, D.M., Doran, T.J., Sinclair, A.H., and Smith, C.A. (2013). Overexpression of aromatase alone is sufficient for ovarian development in genetically male chicken embryos. PLOS ONE 8, e68362. 10.1371/journal.pone.0068362.

81. Fridolfsson, A.K., and Ellegren, H. (1999). A simple and universal method for molecular sexing of non-ratite birds. J. Avian Biol., 116–121.

82. Jalabert, C., Liu, M.Q., and Soma, K.K. (2025). Estrogen profiling in blood and brain: Effects of season and an aggressive interaction in a songbird. PLOS ONE 20, e0326727. 10.1371/journal.pone.0326727.

83. Keski-Rahkonen, P., Desai, R., Jimenez, M., Harwood, D.T., and Handelsman, D.J. (2015). Measurement of estradiol in human serum by LC-MS/MS using a novel estrogen-specific derivatization reagent. Anal. Chem. 87, 7180–7186. 10.1021/acs.analchem.5b01042.

84. Jalabert, C., Shock, M.A., Ma, C., Bootsma, T.J., Liu, M.Q., and Soma, K.K. (2022). Ultrasensitive quantification of multiple estrogens in songbird blood and microdissected brain by LC-MS/MS. eNeuro 9. 10.1523/ENEURO.0037-22.2022.

85. Puelles, L., Martinez-de-la-Torre, M., Paxinos, G., Watson, C., and Martínez, S. (2007). The chick brain in stereotaxic coordinates: an atlas featuring neuromeric subdivisions and mammalian homologies. (No Title), lxxv–97.

86. Andrews, S. (2010). FastQC: a quality control tool for high throughput sequence data. http://www.bioinformatics.babraham.ac.uk/projects/fastqc.

87. Kim, D., Paggi, J.M., Park, C., Bennett, C., and Salzberg, S.L. (2019). Graph-based genome alignment and genotyping with HISAT2 and HISAT-genotype. Nat. Biotechnol. 37, 907–915. 10.1038/s41587-019-0201-4.

88. Okonechnikov, K., Conesa, A., and García-Alcalde, F. (2016). Qualimap 2: advanced multi-sample quality control for high-throughput sequencing data. Bioinformatics 32, 292–294. 10.1093/bioinformatics/btv566.

89. Pertea, M., Pertea, G.M., Antonescu, C.M., Chang, T.-C., Mendell, J.T., and Salzberg, S.L. (2015). StringTie enables improved reconstruction of a transcriptome from RNA-seq reads. Nat Biotechnol 33, 290–295. 10.1038/nbt.3122.

90. Love, M.I., Huber, W., and Anders, S. (2014). Moderated estimation of fold change and dispersion for RNA-seq data with DESeq2. Genome Biol. 15, 550. 10.1186/s13059-014-0550-8.

91. Martínez-Camblor, P., Pérez-Fernández, S., and Díaz-Coto, S. (2020). The role of the p-value in the multitesting problem. J. Appl. Stat. 47, 1529–1542. 10.1080/02664763.2019.1682128.

92. Langfelder, P., and Horvath, S. (2008). WGCNA: an R package for weighted correlation network analysis. BMC Bioinform. 9, 559. 10.1186/1471-2105-9-559.

