## Supplementary figures and images for "Gene expression in the hypothalamus of gonad-sex chromosome divergent embryonic chickens"

### Supplementary Figure 1

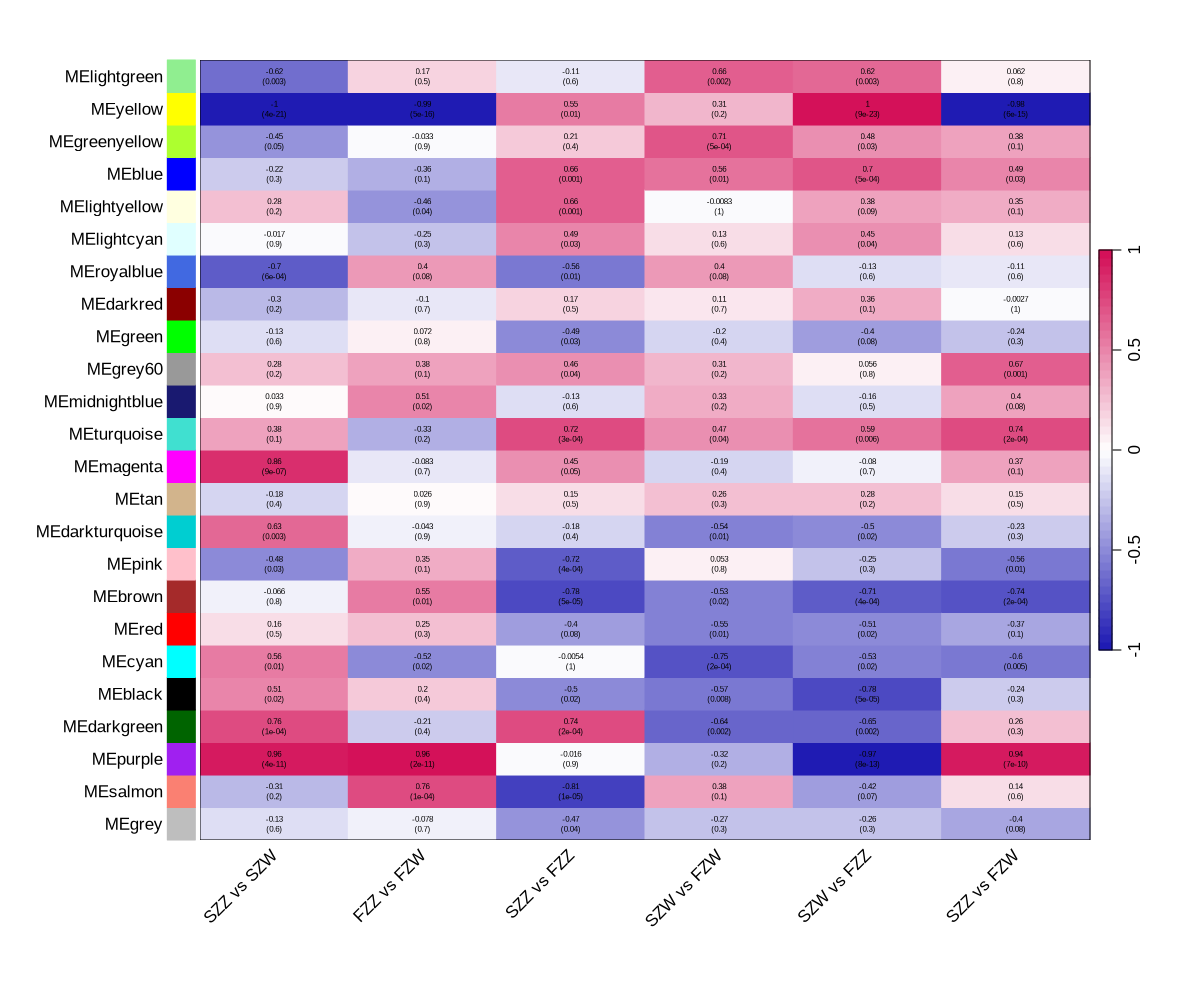

### Supplementary Figure 2

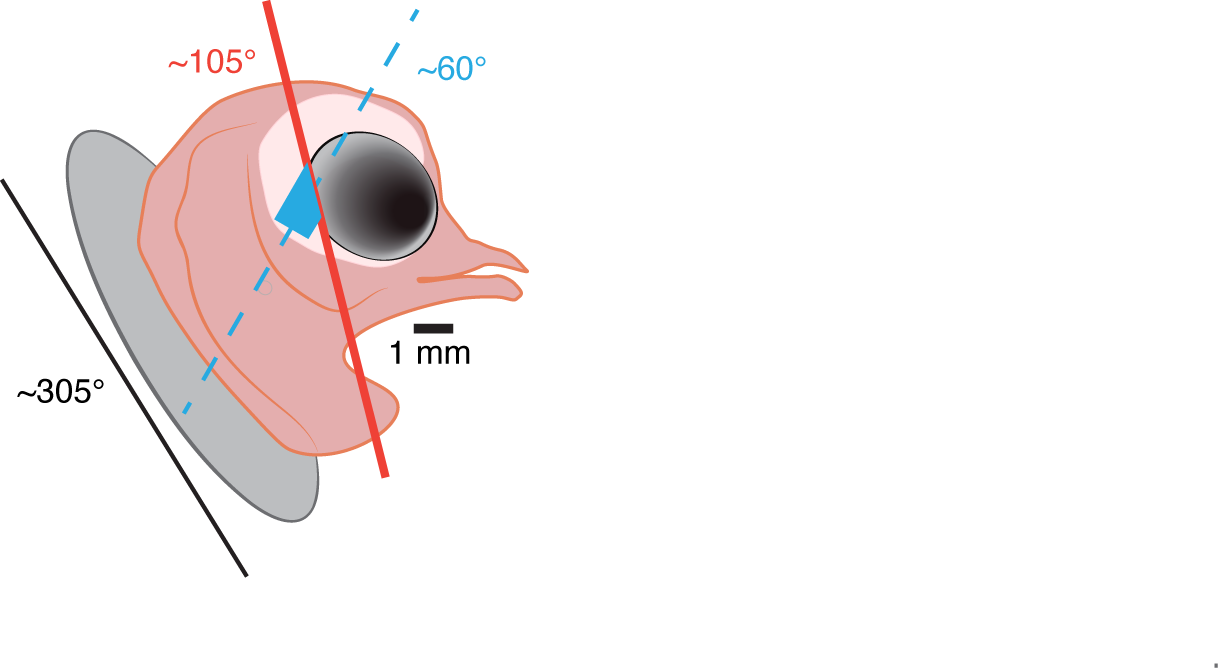
